# Evidence for multiple independent expansions of Fox gene families within flatworms

**DOI:** 10.1101/2025.01.09.632129

**Authors:** Ludwik Gąsiorowski

## Abstract

Expansion and losses of gene families are important drivers of molecular evolution. A recent survey of Fox genes in flatworms revealed that this superfamily of multifunctional transcription factors, present in all animals, underwent extensive losses and expansions during platyhelminth evolution. In this paper, I analyzed Fox gene complement in four additional species of platyhelminths, that represent early-branching lineages in the flatworm phylogeny: catenulids (*Stenostomum brevipharyngium* and *Stenostomum leucops*) and macrostomorphs (*Macrostomum hystrix* and *Macrostomum cliftonense*). Phylogenetic analysis of Fox genes from this expanded set of species provided evidence for multiple independent expansions of Fox gene families within flatworms. Notably, *FoxG*, a panbilaterian brain-patterning gene, appears to be the least susceptible to duplication, while *FoxJ1*, a conserved ciliogenesis factor, has undergone extensive expansion in various flatworm lineages. Analysis of the single-cell atlas of S. *brevipharyngium*, combined with RNA *in situ* hybridization, elucidated the tissue-specific expression of the selected Fox genes: *FoxG* is expressed in the brain, three of the Fox genes (*FoxN2/3-2, FoxO4* and *FoxP1*) are expressed in the pharyngeal cells of likely glandular function, while one of the *FoxQD* paralogs is specifically expressed in the protonephridium. Overall, the evolution of Fox genes in flatworms appears to be characterized by an early contraction of the gene complement, followed by lineage-specific expansions that have enabled the co-option of newly evolved paralogs into novel physiological and developmental functions.

**Statements and Declarations:** The author has no competing interests to declare that are relevant to the content of this article. The research was funded by the Alexander von Humboldt Foundation (The Humboldt Research Fellowship for Postdoctoral Researchers) and The Polish National Agency for Academic Exchange (Polish Returns NAWA grant no. BPN/PPO/2023/1/00002).

## Introduction

Expansions and losses of gene families are among the main driving forces of molecular evolution (Albalat and Cañestro 2016; Fernández and Gabaldón 2020; Lespinet et al. 2002; Susumu 1970). The expansions, resulting from the duplications of a single ancestral gene, are often followed by the gain of new functions for one or more of its descendant copies, which allow the evolution of new phenotypes (Kryuchkova-Mostacci and Robinson-Rechavi 2016; Langleib et al. 2024; Lespinet et al. 2002; Mantica et al. 2024; Susumu 1970). On the other hand, losses of the already existing genes might be associated with morphological or physiological simplification of the organism, which under certain conditions (e.g., body size reduction, life cycle simplification, parasitism), can also show adaptive value (Gregory et al. 2000; Gross et al. 2019; Liao et al. 2023; Malmstrøm et al. 2018; Slyusarev et al. 2020). Additionally, both gene losses and duplication might be also attributed to the large-scale changes in the genome (such as whole genome duplications and chromosomal rearrangements) that are considered important drivers of genome evolution (Lewin et al. 2024; Liao et al. 2023; Plessy et al. 2024). Altogether, it is evident that the evolutionary history of the organisms will have a strong imprint on the complement and content of the conserved gene families present in its genome.

Fox genes (also known as forkhead box genes) encode for a highly conserved superfamily of transcription factors, involved in multiple developmental and physiological processes (Carlsson and Mahlapuu 2002; Hannenhalli and Kaestner 2009). The number and complement of Fox genes in particular organisms were shaped by the combination of duplication and losses that occurred since the ancestral proto-Fox originated in the common ancestor of animals and fungi (Larroux et al. 2008; Mazet et al. 2003; Shimeld et al. 2010). The rapid expansion of Fox genes at the base of animal phylogeny resulted in the emergence of 23 Fox families (named with the consecutive letters of the Latin alphabet) in the last common ancestor of Bilateria (Fritzenwanker et al. 2014; Mazet et al. 2003; Pascual-Carreras et al. 2021; Seudre et al. 2022; Shimeld et al. 2010). Since then, the Fox complement has been conserved in some animal lineages (for instance 19 of those are still present as single copies in annelids *Owenia fusiformis* and *Capitella teleta* (Seudre et al. 2022) as well as in hemichordate *Saccoglossus kowalevskii* (Fritzenwanker et al. 2014) and Onychophoran *Euperipatoides kanangrensis* (Janssen et al. 2022)) while in others, multiple Fox families have been lost altogether (for example tardigrades preserved only 13 and nematodes 12 of the ancestral families (Janssen et al. 2022)).

Flatworms (Platyhleminthes) are among the animal clades, in which the evolution of Fox genes was particularly dynamic. The recent survey of Fox gene complement in different groups of flatworms (Pascual-Carreras et al. 2021) has shown massive loss of the conserved bilaterian Fox genes in the common ancestor of all plathelminths, followed by expansions of the remaining families. While this work incorporated information on the Fox complement from a broad diversity of flatworms, whose transcriptomes or genomes were available at the time of publication, it suffered from the limited availability of high-quality transcriptomes and genomes of two first offshoots of platyhelminth phylogeny – catenulids and macrostomorphs (Egger et al. 2015; Larsson and Jondelius 2008; Laumer et al. 2015). Both clades were represented only by a single species – the first one by a transcriptome of *Stenostomum sthenum* (referred to as “Catenulida” in the paper), and the latter by a genome of *Macrostomum lignano*. Especially, the inclusion of *M. lignano* as a solely representative of Macrostomorpha can be problematic, as *M. lignano* went through recent genome duplication (Zadesenets et al. 2017a; Zadesenets et al. 2017b) that hampers the reconstruction of the ancestral macrostomids conditions and thus hazes the image of the evolution of their gene content.

To overcome these limitations, I took advantage of the recent publications of high-quality transcriptomes of additional *Stenostomum* species (Gąsiorowski et al. 2023a) and genomes of two species of *Macrostomum* that branched off the *M. lignano* lineage before its genome duplication occurred (Wiberg et al. 2023). The survey of Fox gene complement in this expanded set of flatworms allows a more precise reconstruction of Fox gene complements within the platyhelminths. Importantly, with the use of the phylogenetic approach, it is now possible to test whether expansions of the limited Fox families retained by the last common flatworm ancestor occurred early on in the flatworm evolution or whether they expanded many times independently in different groups of flatworms. Finally, the availability of the single-nuclei transcriptome of the catenulid *S. brevipharyngium* (Gąsiorowski et al. 2024) and the possibility of performing RNA *in situ* hybridization in this species (Gąsiorowski et al. 2023a) allows insight into the spatial and cell type-specific expression of its Fox genes, which sheds light on the possible functions of the particular Fox genes in this species and how they evolve within flatworms.

## Material and methods

### Phylogenetic analyses

First, I searched for sequences encoding putative Fox genes in the published transcriptomes of two species of *Stenostomum, S. brevipharyngium* (PRJNA1004231 published in Gąsiorowski et al. (2023a)) and *S. lecuops* (PRJNA276469 published in Laumer et al. (2015)) and genomes of two species of Macrostomum, *M. cliftonense* and *M. hystrix* published in Wiberg et al. (2023). The reference Fox protein sequence (FoxJ1 from *Saccoglossus kowalevskii*, NP_001158438.1) was used to perform tblastn search in the assembled non-redundant transcriptomes (for *Stenostomum*) and collection of the longest transcripts assembled to the genomes (for *Macrostomum*). The nucleotide sequences obtained with this initial search were blasted back (with a blastx algorithm) against protein sequence collection at NCBI. Only sequences that gave reciprocal blast hits against other metazoan Fox genes were kept in the following analyses (Online Resource 1: Table S1). Sequences of all newly identified genes have been deposited in GenBank (accession numbers PQ406716 – PQ406829).

To assign the obtained sequences into metazoan Fox families, I realign them with an alignment of metazoan Fox genes published in Fritzenwanker et al. (2014) and additional reference sequences (Online Resource 1: Table S2) using Clustal Omega (v 1.2.3) (Sievers and Higgins 2014) implemented in Geneious Prime (v2023.0.3). The alignment was trimmed in Geneious Prime to remove sites containing more than 50% gaps before being used for phylogenetic analyses. This full Fox alignment was then analyzed with FastTree (v 2.1.11) (Price et al. 2010) implemented in Geneious Prime and raxmlGUI (v 2.0) (Edler et al. 2021) to generate phylogenetic trees of all Fox genes. The LG amino acid substitution model was used for both analyses based on the results of ModelTest-NG v0.1.7 (Darriba et al. 2020) implemented in raxmlGUI.

To analyze the evolution of particular Fox families within flatworms, I aligned the newly obtained sequences with Fox sequences from the same Fox family in other flatworms originating from the analysis of Pascual-Carreras et al. (2021) (Online Resource 2). The alignments for each Fox family were trimmed in Geneious Prime to remove sites containing more than 50% gaps and analyzed with raxmlGUI. For each analysis, I used the best-scored model of amino acid substitution as indicated by ModelTest-NG v0.1.7 implemented in raxmlGUI (Online Resource 1: Table S3). To reconstruct duplications of ancestral Fox genes in flatworms I manually analyzed the obtained phylogenies of Fox families, searching for patterns of well-supported sister clades, each containing identical or similar sets of species.

### Analysis of cell type-specific expression

For analyzing the specificity of Fox gene expression to particular cell types of *S. brevipharyngium*, I made use of its recently published single-cell atlas (Gąsiorowski et al. 2024). A dotplot was generated by calculating the percentage of cells expressing each gene and the mean expression levels within each cell cluster from the SoupX-corrected single-nuclei RNA-seq data (PRJNA1156255). Expression values were extracted using Scanpy (Wolf et al. 2018), and percentages were computed by dividing the number of cells expressing a gene by the total number of cells in each cluster. Both variables were visualized using Matplotlib.

### In situ hybridization

For animal husbandry, fixation, and hybridization chain reaction v 3.0 (Choi et al. 2018), I followed the same protocol as in (Gąsiorowski et al. 2023a). The HCR probe sets were designed using Özpolat Lab’s HCR in situ probe generator (Kuehn et al. 2022). Pools of HCR probes are available in Online Resource 1: Table S4. Stained animals were imaged on an Olympus IX83 microscope with a spinning disc Yokogawa CSUW1-T2S scan head, and the obtained confocal Z-stacks were processed for contrast and brightness in Fiji (Schindelin et al. 2012).

## Results

### Fox complement in *Stenostomum* and *Macrostomum*

The search for Fox genes resulted in the identification of 25 putative Fox genes in *S. brevipharyngium*, 23 genes in *S. leucops*, 32 genes in *M. cliftonense* and 34 genes in *M. hystrix* (Online Resource 1: Table S1). Phylogenetic analyses of metazoan Fox genes (Fig. 1 a and b; Online Resource 3: Figures S1 and S2) allowed the assignment of most of those genes into well-conserved Fox families (Fig 1c). In terms of the presence or absence of the ancestral metazoan fox families in catenulids and macrostomorphs my results are similar to those of Pascual-Carreras et al. (2021). In all four analyzed species, I failed to find representatives of *FoxB, FoxAB, FoxE, FoxH, FoxI, FoxL2, FoxM, FoxN1/4, FoxQ1, FoxQ2* and *FoxT* families (Fig. 1c). Additionally, all stenostomids seems to be lacking *FoxJ1*. In contrast to the previous findings, I identified a single member of *FoxC* family in one of the analyzed catenulids, *S. brevipharyngium*, that was missing from the transcriptomes of both *S. sthenum* and *S. leucops*. Also, I retrieved two paralogs of *FoxJ1* from both newly analyzed *Macrostomum* species, providing evidence of the existence of this Fox family in macrostomorphs, in which it has been previously reported as missing. Non-canonical Fox genes, that could not be placed within well-defined Fox clades, were present both in *Stenostomum* and *Macrostomum* but only in one or two copies per species. Among those two of the non-canonical Fox genes from *S. leucops* (ex_Sleu_v1_3482_1_1 and ex_Sleu_v1_63544_1_1) were giving sequences of Fox genes from Microsporidia as best BLAST hits (Online Resource 1: Table S5). In both analyzed genera, the number of paralogs of particular Fox genes was species-specific showing a variation from single copies (for instance *FoxG* in all three species of *Stenostomum*) to 8 copies of *FoxC* in *M. lignano*.

**Fig. 1.**
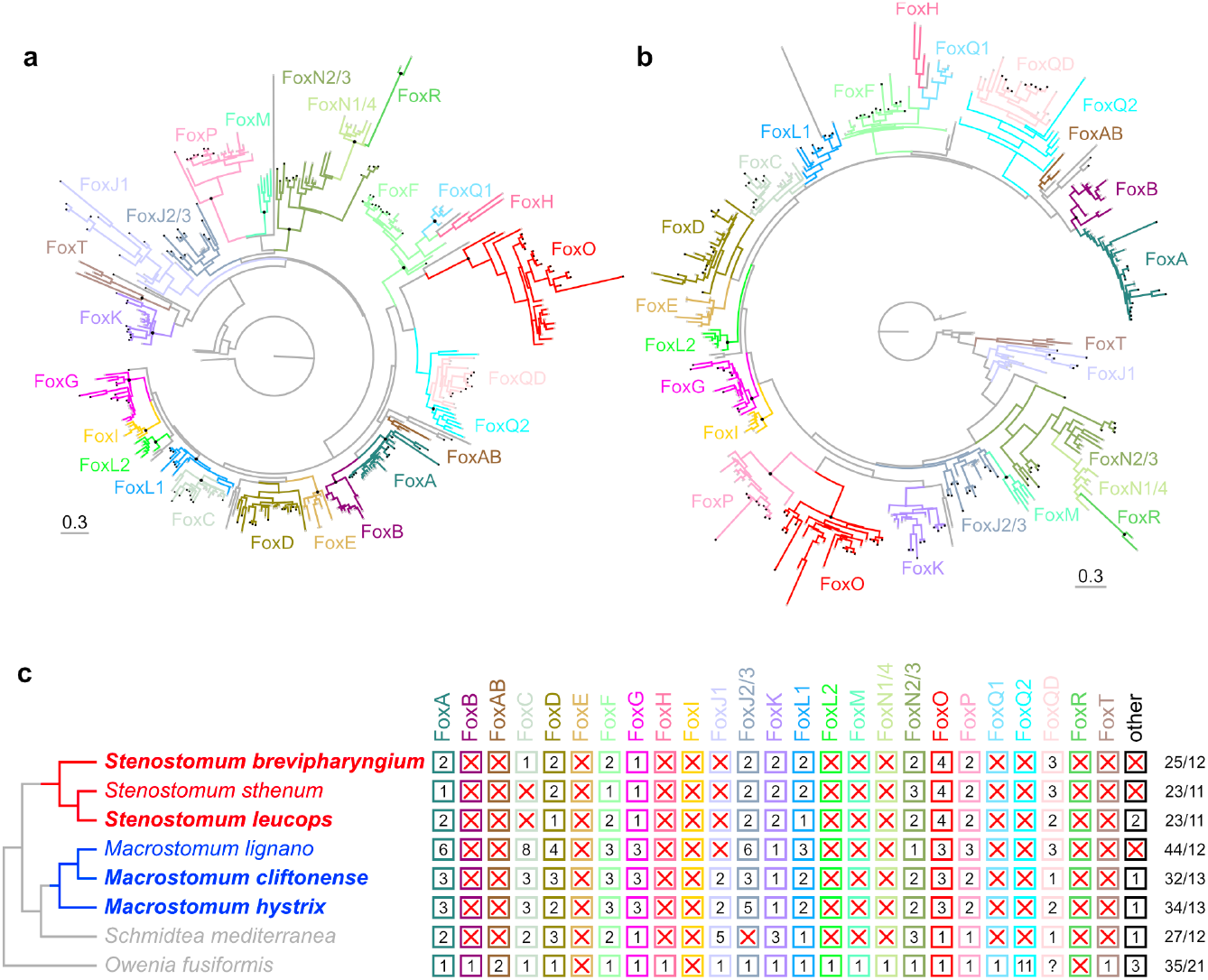
Complement of Fox genes in *Stenostomum* and *Macrostomum* based on phylogenetic analysis of candidate sequences. **a**. Phylogeny of metazoan Fox genes, approximately-maximum-likelihood tree computed with FastTree (v 2.1.11) under LG amino acid substitution model. **b**. Phylogeny of metazoan Fox genes, maximum-likelihood tree computed with raxmlGUI (v 2.0) under LG amino acid substitution model. For both **a** and **b**, black squares at terminals indicate sequences of flatworms, and black dots at nodes indicate bootstrap support greater than 90%. The trees with terminal names and exact support values can be found as Figures S1 and S2 in Online Resource 3. **c**. Complement of Fox genes in representatives of *Stenostomum* and *Macrostomum*, compared to a planarian *Schmidtea mediterranea* and an annelid *Owenia fusiformis*. Species in bold were analyzed in this study. The complement for *M. lignano, S. sthenum* and *S. mediterranea* comes from the analysis of Pascual-Carreras et al. (2021) and complement for *O. fusiformis* from Seudre et al. (2022). The last column provides the number of Fox sequences and the number of conserved bilaterian Fox families in each species.

### Analysis of Fox gene duplications

To gain insight into whether multiple copies of particular Fox genes present across flatworm phylogeny originate from ancient or more recent expansions of ancestral genes, I performed phylogenetic analyses focused on flatworm sequences. Analysis of the phylogeny of particular Fox gene families in flatworms (Online Resource 3: Figs. S3-S15) allowed the identification of putative duplication events (Fig. 2a; see Material and methods for details). The number of reconstructed duplications was unevenly distributed among analyzed clades, with a decisive majority of them occurring closer to the leaves of the tree – 18 in the genus *Macrostomum*, 17 additional ones in the lineage of *M. lignano*, and 12 in the lineage of *Stenostomum* (Fig. 2a and b). At the same time, in the common ancestor of Rhabditophora only a single duplication of *FoxJ1* was reconstructed, and none of the duplications was reconstructed in the last common ancestor of all flatworms (Fig. 2a and b). I also scored the number of reconstructed duplications by the Fox gene family (Fig. 2c). While most of the families showed between 4 and 6 duplication events among 15 analyzed flatworm species, there were some apparent outliers. In particular, there was only one clear duplication of *FoxG* in the common ancestor of the three analyzed *Macrostomum* species, while *FoxJ1* showed 12 putative duplication events at different phylogenetic levels, from ancient one (in the common ancestor of Rhabditophora), through intermediate ones (e.g., in the lineage of *Macrostomum* spp., or the common ancestor of planarians) to more recent ones (for example in the lineage of *Dendrocoelum lacteum*). Overall, the analysis of Fox gene duplications in flatworms suggests that while duplications are widespread, they occurred independently across different lineages, are mostly limited to more recent clades, and exhibit distinct clade- and gene family-specific duplication patterns.

**Fig. 2.**
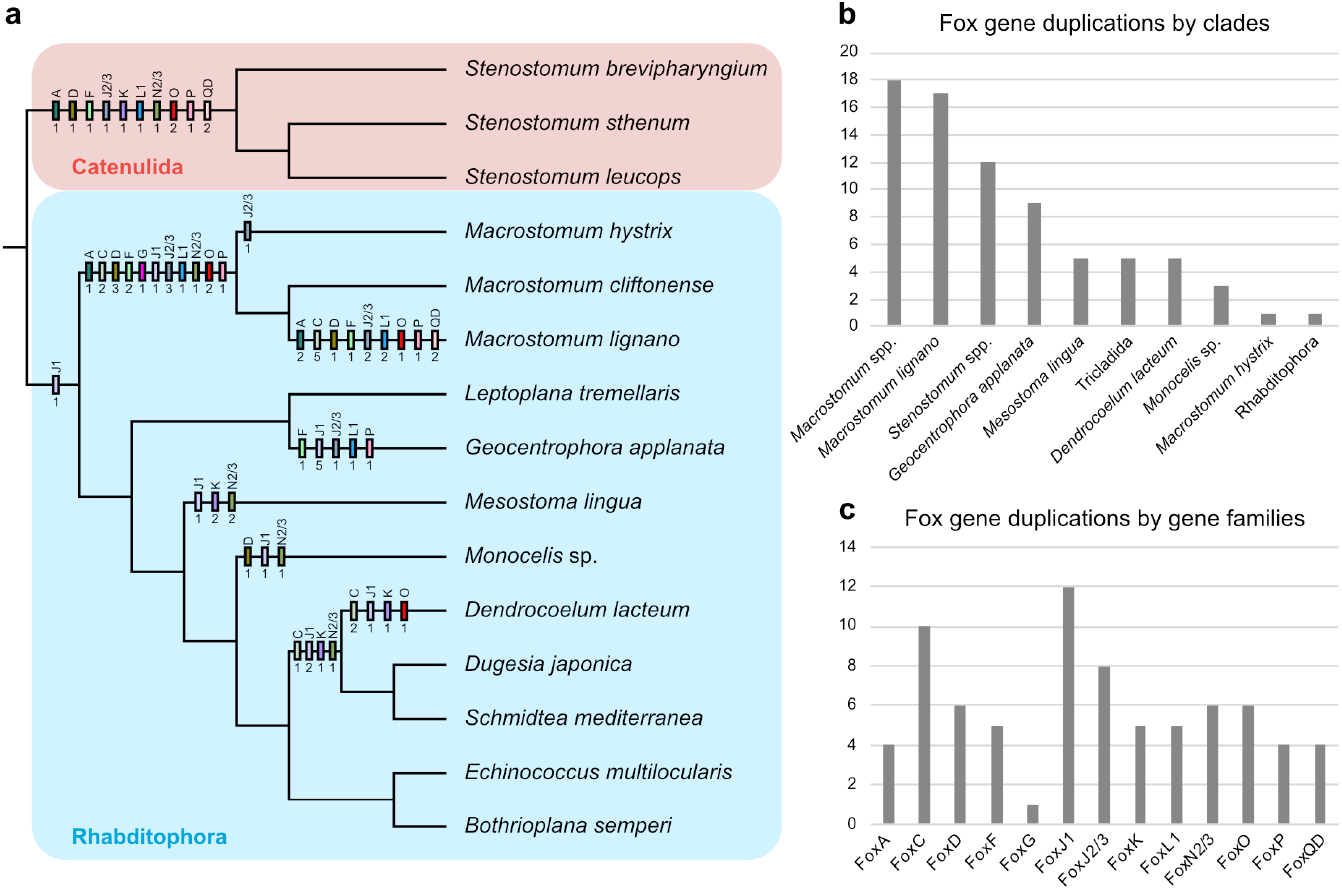
Results of the analysis of Fox gene duplications in flatworms. **a**. Phylogenetic distribution of reconstructed Fox duplications on flatworm phylogeny. Each rectangle represents duplications of a particular Fox gene on a given branch. The name of the duplicated Fox gene is provided above the rectangle and the number of reconstructed duplications of the given gene is below. Tree topology follows Laumer et al. (2015). **b**. The number of reconstructed Fox gene duplications in different clades of flatworms. **c**. The number of reconstructed duplications of particular Fox families in flatworms.

### Expression in single-cell atlas

Analysis of the single-cell atlas revealed a varying level of expression and specificity of Fox genes to particular cell types of catenulid *S. brevipharyngium* (Fig. 3). Some of the Fox genes are barely expressed in the dataset (for example, *FoxC, FoxQD-2, FoxQD-3*), while others show relatively uniform expression across most cell types (*FoxF1, FoxN2/3-1*). Few of the analyzed genes exhibit cell-type specific expression – *FoxG* in clusters “nerve cord neurons”, and “neurons I and muscles”; *FoxN2/3-2* and *FoxO4* in cluster “pharyngeal cells I”, *FoxP1* in “brain neurons” and to lesser extent in “pharyngeal cells I” and *FoxQD-1* in “protonephridial cells”.

**Fig. 3.**
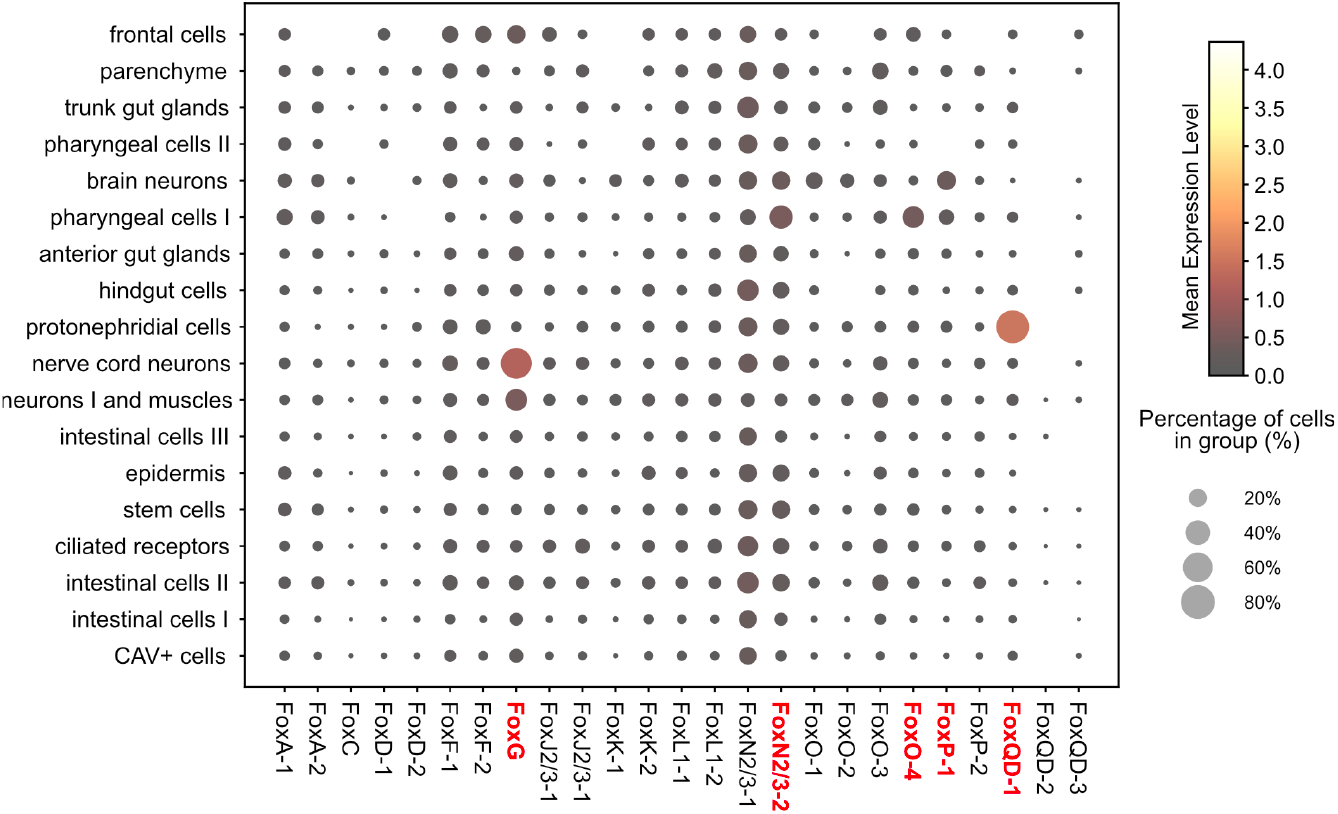
Dotplot showing the specificity of expression of particular Fox genes in different cell types of *Stenostomum brevipharyngium*. Cell types names follow the nomenclature from the original report of a single-cell atlas (Gąsiorowski et al. 2024). Genes, which expression patterns were investigated with *in situ* RNA hybridization, are indicated in red.

### In situ hybridization of selected Fox genes

Based on the analysis of the single-cell atlas I selected five Fox genes that show cell-type specific expression patterns for *in situ* RNA hybridization – *FoxG, FoxN2/3-2, FoxO4, FoxP1* and *FoxQD-1*. Each of those genes was hybridized together with molecular markers of the cell types in which it showed elevated expression (based on Gąsiorowski et al. (2024)), to confirm the cell type specificity of particular Fox genes. In general RNA *in situ* hybridization showed similar results as *in silico* analysis of cell type-specific Fox expression. *FoxG* is expressed in posterior brain neurons, where it co-localizes with *UNC13* and *SYT16*, two neuronal markers (Fig. 4a). However, I was not able to detect *FoxG*^+^ cells in the longitudinal nerve cords, in which *SYT16* is highly expressed. Both *FoxN2/3-2* and *FoxO4* are expressed in the *RBM24*^+^ large pharyngeal cells located in the dorso-anterior pharynx (Fig. 4b and c). Additionally, *FoxN2/3-2* is also expressed in the more posterior part of the pharynx (Fig. 4b), while *FoxO4* is expressed in the dorsal part of the head, in front of the pharynx, likely in the most posterior part of the dorsal brain lobes (Fig. 4c). Similarly, *FoxP1* is expressed in the *MEF2C*^+^ brain neurons, but also in the *RBM24*^+^ pharyngeal cells (Fig. 4d). Finally, *FoxQD-1*, is highly, and specifically expressed in the protonephridium, as indicated by single-cell analysis, where it is strictly co-expressed with the protonephridial marker SLC4A7 in a zigzagging dorsal domain (Fig. 4e).

**Fig. 4.**
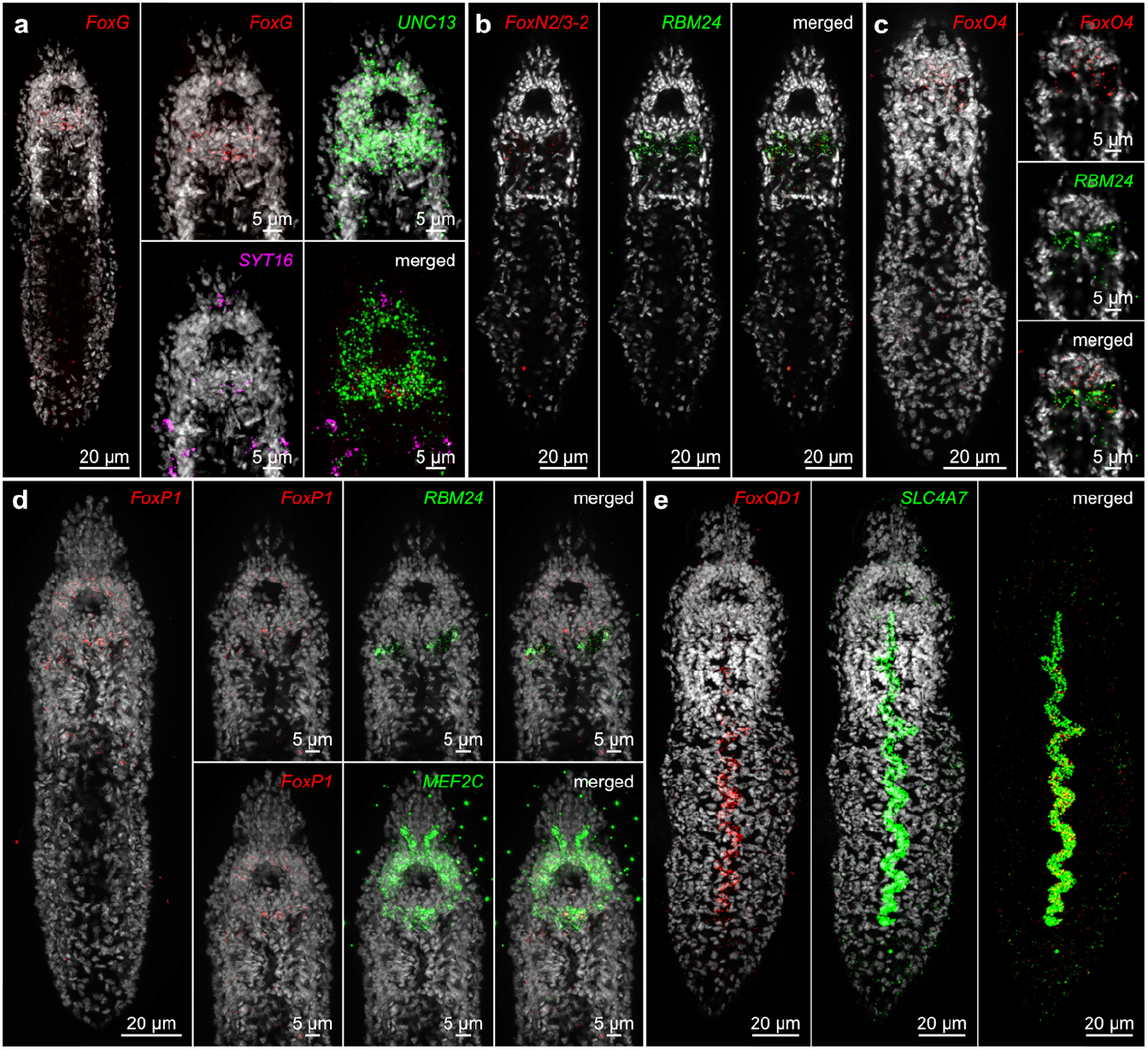
RNA *in situ* hybridization of selected Fox genes and cell type markers in *Stenostomum brevipharyngium*. **a**. *FoxG* is expressed in the posterior brain, where it is co-expressed with the neuronal markers *UNC13* and *SYT16*. **b**. *FoxN2/3-2* is expressed in the *RBM24*^*+*^ cells in the dorso-anterior pharynx. **c**. *FoxO4* is expressed in the *RBM24*^*+*^ cells in the dorso-anterior pharynx and more anteriorly in the dorso-posterior brain. **d**. *FoxP1* is expressed throughout the brain, where it is co-expressed with the brain neuron marker *MEF2C* and in the *RBM24*^*+*^ cells in the dorso-anterior pharynx. **e**. *FoxQD1* is co-expressed with the protonephridial marker *SLC4A7* in the dorsally located excretory organ.

## Discussion

### Dynamic evolution of Fox genes in flatworms

The Fox gene complement has now been studied in multiple species of flatworms from all major evolutionary lineages (Pascual-Carreras et al. 2021), allowing a comprehensive reconstruction of the evolution of Fox genes within this animal clade. The systematic absence of several Fox families, otherwise conserved in Metazoa (Fritzenwanker et al. 2014; Janssen et al. 2022; Larroux et al. 2008; Pascual-Carreras et al. 2021; Seudre et al. 2022; Shimeld et al. 2010), in all tested flatworm species, indicates a massive ancestral loss of Fox gene in the last common flatworm ancestor. With the loss of 11 ancestral metazoan Fox families (*FoxB, FoxAB, FoxE, FoxH, FoxI, FoxL2, FoxM, FoxN1/4, FoxQ1, FoxQ2* and *FoxT*), the ancestral flatworm Fox complement consisted of only 12 families. Yet, most of the flatworms have between 15 to 35 Fox genes, indicative of subsequent expansions of the remaining Fox genes. The results of analyses performed in this study show that most of those duplications can be attributed to relatively recent nodes in flatworm phylogeny, indicating multiple expansions of Fox genes that occurred independently in different clades of flatworms. In line with previous reports of recent rounds of whole genome duplication in *Macrostomum lignano* (Zadesenets et al. 2017a; Zadesenets et al. 2017b), I found evidence of multiple duplications of particular Fox genes that occurred only in the lineage of *M. lignano*, but are missing in other *Macrostomum*, and which contributed to the record number of 44 Fox genes in this species. Interestingly, none of the studied flatworms preserved the ancestral platyhelminth Fox complement, and even those species that have relatively few Fox genes (for instance polyclad *Leptoplana tremellaris* or bothrioplanid *Bothrioplana semperi* both with only 10 Fox genes (Pascual-Carreras et al. 2021)) have multiple copies of some Fox genes and lost Fox families that were present in the common flatworm ancestor. All in all, the overall evolution of Fox complement in flatworms seems to be marked by very dynamic patterns of gene losses and duplications.

It is important to mention here, that similar patterns of losses and subsequent independent re-expansions are also present in other conserved gene families studied in flatworms, for example, for Wnt and Hox genes (Currie et al. 2016; Gąsiorowski et al. 2023b; Liao et al. 2023; Riddiford and Olson 2011). This might indicate that the common ancestor of all flatworms went through extensive genome reduction, which resulted in massive losses of many genes that remain conserved among other metazoans. Such extensive reduction of the ancestral genome could also partially explain a recently reported lack of metazoan ancestral linkage groups in the genomes of multiple flatworm species (Ivanković et al. 2024). The putative cause of this hypothetical genomic reduction remains a mystery. One possibility is that it was related to body miniaturization – microscopic flatworms form a basal grade within flatworm phylogeny (Egger et al. 2015; Larsson and Jondelius 2008; Laumer et al. 2015), indicating that the last common ancestor of all flatworms probably had a relatively small body plan. Reduction and reshuffling of the genome in the instance of miniaturization is a common mechanism that has been reported e.g., in annelids (Martín-Durán et al. 2021), or tardigrades (Gross et al. 2019). On the other hand, even in microscopic flatworm lineages, for example, catenulids and macrostomorphs, the reduced set of Fox genes was not retained and it re-expanded, indicating that the miniature body plan of flatworms is not as strictly linked with the reduced absolute number of genes.

### Conservation of FoxG across flatworms

*FoxG* was identified in this analysis as the least susceptible to duplications among all Fox genes present in flatworms, suggesting that this gene might be more strongly conserved. *FoxG*, also known as *brain factor 1 (BF1)*, has a conserved function in the patterning of anterior neuronal domains across Bilateria (Kumamoto and Hanashima 2017; Luo et al. 2018; Toresson et al. 1998). Although in planarians, its expression in the brain is not very specific (Koinuma et al. 2003; Pascual-Carreras et al. 2021), in *Stenostomum*, the gene has an exclusive expression domain in the brain (this study). The involvement of *FoxG* in the patterning of the brain, a structure that is crucial for most free-living bilaterians regardless of their size, might explain a relative conservation of this particular Fox gene in flatworms. Interestingly, *FoxG* was duplicated in a planarian *Polycelis nigra* (two copies (Pascual-Carreras et al. 2021)), and all studied macrostomorphs. Future studies on the expression and function of *FoxG* paralogs in those species could hint at whether those newly evolved paralogs evolved new roles or just maintained a redundancy in brain patterning mechanisms.

### Lability of FoxJ1

In contrast to *FoxG, FoxJ1* showed the most dynamic evolutionary history among flatworms, exemplified by comparatively long branches in phylogenetic trees, as well as complex patterns of lineage-specific losses and expansions. Although *FoxJ1* has been reported as missing in both catenulids and macrostomorphs by Pascual-Carreras et al. (2021), here I was able to identify two paralogs of *FoxJ1* in two species of *Marcostomum*, while confirming the apparent lack of the gene in all tested stenostomids. Remarkably, this pattern of presence and absence makes *FoxJ1* the only Fox gene showing ancient divergence within flatworms – seemingly it was lost in the catenulid (or at least *Stenostomum*) lineage and duplicated in the ancestor of Rhabditophora, followed by multiple additional rounds of expansions in various rhabditophoran clades.

*FoxJ1* has a known conserved function in the ciliogenesis of motile cilia (Cruz et al. 2010; Stubbs et al. 2008; Vij et al. 2012; Yu et al. 2008). Flatworms heavily rely on motile ciliation for locomotion, feeding and excretion (Jennings 1957; Rohde 1991; Tyler 1984; Vu et al. 2019; Vu et al. 2015). Therefore, duplicated copies of *FoxJ1* in particular lineages might be related to the subfunctionalization of *FoxJ1* paralogs into the different types of ciliated cells. For instance, in *Schmidtea mediterrana*, particular *FoxJ* paralogs show divergent expression domains: *FoxJ1-4* is expressed in the ventral ciliated epidermal cells and dorsal mid-line ciliated cells, while *FoxJ1-1* and FoxJ1-2 only in ciliated cells of the dorsal midline (Vij et al. 2012). The systematic lack of *FoxJ1* in all studied *Stenostomum* species is perplexing, as catenulids are heavily ciliated and use motile cilia for locomotion, excretion and food processing (Jennings 1957; Rohde and Watson 1993; Tyler 1984). Future studies are needed to explore how this intricate ciliation functions without one of the core ciliogenetic factors.

### Expansions of FoxO

*FoxO* has a conserved function in controlling cell death, also maintained in planarians (Pascual-Carreras et al. 2021). The previous survey of Fox genes in flatworms showed that while most platyhelminths have only one or two copies of *FoxO*, multiple copies exist in *Stenostomum sthenum* (4) and in *Macrostomum lignano* (3) (Pascual-Carreras et al. 2021). My analysis of additional species of *Macrostomum* and *Stenostomum* reveals that multiple copies of *FoxO* in those species originated by a series of duplications that occurred before the split of analyzed species, independently in catenulids and macrostomorphs. Although we lack information on the function of these *FoxO* paralogs in each of the species, the analysis of *FoxO* expression among different cell types of *S. brevipharyngium* revealed that particular paralogs are expressed in different cell types, indicating functional divergences of this otherwise well-conserved gene in stenostomids.

### Fox genes in Catenulida

The inclusion of additional species of *Stenostomum* and the possibility of investigation of gene expression with single cell-atlas and *in situ* hybridization for *S. brevipharyngium* allow a deeper understanding of Fox genes evolution and function in catenulids. For instance, each of the transcriptomes of two additional species of *Stenostomum* that were analyzed in this study contained two paralogs of *FoxA* and *FoxF*, while both of those genes were reported as unduplicated in *S. sthenum* (Pascual-Carreras et al. 2021). Moreover, the transcriptome of *S. brevipharyngium* contained a single paralog of *FoxC*, providing the first evidence of the presence of this particular Fox family in any catenulid species. Interestingly, two of the non-canonical Fox genes from *S. leucops* show strong equences similarity to Fox genes from Microsporidia, unicellular intracellular parasitic fungi. As Microsporidia are common parasites of aquatic invertebrates, including flatworms (Stentiford and Dunn 2014), it might indicate that the animals used to generate the transcriptome were infested with the microsporidial symbiont or that the genes were horizontally transferred to the host.

The analysis of the single-cell atlas provided the first glance into the expression of particular Fox genes in *S. brevipharyngium*. While most did not show strong cell type specificity, there were a few exceptions, which I could confirm with the RNA *in situ* hybridization. Unsurprisingly, *FoxG* is specifically expressed in the brain neurons, in line with its conserved function in brain patterning across Bilateria (Kumamoto and Hanashima 2017; Luo et al. 2018; Toresson et al. 1998). Three of the Fox genes, *FoxN2/3-2, FoxO4* and *FoxP1* were expressed in the “pharyngeal cells I”. This cell type, of yet unknown function, has been recently characterized based on single-cell transcriptomic data (Gąsiorowski et al. 2024). Those large cells, located in the dorso-anetior pharynx, express *RNA-binding protein 24-A-like* (*RBM24*) along with batteries of *Stenostomum*-specific genes that remain functionally uncharacterized. Taking into account their anatomical location, morphology and expressional profile, the “pharyngeal cells I” likely represent glandular cells associated with the anterior section of the digestive system. As for now, homologous cell types are unknown from the other flatworms. Therefore, the expression of 3 Fox genes that originated through Stenostomum-specific gene duplications in this *Stenostomum*-specific cell type that expresses *Stenostomum*-specific genes, represents an elegant example of neofunctionalization of recently duplicated transcription factors into the patterning of evolutionary novel cell type. Finally, I also found a strikingly specific and strong expression of one of the *Stenostomum*-specific *FoxQD* paralogs in protonephridium, indicating the involvement of this gene in the development or maintenance of the excretory organ. This represents the first report of *FoxQD* expression in excretory organs of any bilaterian and likely it resulted from the recent co-optation of this particular paralog into the patterning of the catenulid excretory system.

## Conclusions

The analysis of Fox gene complement and phylogeny in additional flatworm species reveals ancient losses of Fox genes in the last common plathelminth ancestor and recent, independent expansion of Fox genes in multiple lineages. These events are predominantly confined to relatively recent clades, with minimal evidence of duplications at deeper nodes of the flatworm phylogeny. The patterns of duplication display clade- and gene family-specific biases, suggesting distinct evolutionary pressures possibly related to the function of particular Fox genes. Analysis of expression of Fox genes in catenulid *S. brevipharyngium*, provides evidence for neofunctionalization of some of those recently evolved Fox paralogs.

## Supporting information

FASTA file containing sequences of flatworm Fox proteins compiled from Pascual-Carreras et al. (2021)

## Supplementary Information

- Online Resource 1 – Supplementary Tables S1-S5
- Online Resource 2 – FASTA file containing sequences of flatworm Fox proteins compiled from Pascual-Carreras et al. (2021)
- Online Resource 3 – Supplementary Figures S1-S15

## References

Albalat R, Cañestro C (2016) Evolution by gene loss. Nature Reviews Genetics 17:379

Carlsson P, Mahlapuu M (2002) Forkhead Transcription Factors: Key Players in Development and Metabolism. Developmental Biology 250:1

Choi HMT, Schwarzkopf M, Fornace ME, Acharya A, Artavanis G, Stegmaier J, Cunha A, Pierce NA (2018) Third-generation in situ hybridization chain reaction: multiplexed, quantitative, sensitive, versatile, robust. Development 145

Cruz C, Ribes V, Kutejova E, Cayuso J, Lawson V, Norris D, Stevens J, Davey M, Blight K, Bangs F, Mynett A, Hirst E, Chung R, Balaskas N, Brody SL, Marti E, Briscoe J (2010) Foxj1 regulates floor plate cilia architecture and modifies the response of cells to sonic hedgehog signalling. Development 137:4271

Currie KW, Brown DD, Zhu S, Xu C, Voisin V, Bader GD, Pearson BJ (2016) HOX gene complement and expression in the planarian Schmidtea mediterranea. EvoDevo 7:7

Darriba D, Posada D, Kozlov AM, Stamatakis A, Morel B, Flouri T (2020) ModelTest-NG: a new and scalable tool for the selection of DNA and protein evolutionary models. Molecular biology and evolution 37:291

Edler D, Klein J, Antonelli A, Silvestro D (2021) raxmlGUI 2.0: A graphical interface and toolkit for phylogenetic analyses using RAxML. Methods in Ecology and Evolution 12:373

Egger B, Lapraz F, Tomiczek B, Müller S, Dessimoz C, Girstmair J, Škunca N, Rawlinson KA, Cameron CB, Beli E (2015) A transcriptomic-phylogenomic analysis of the evolutionary relationships of flatworms. Current Biology 25:1347

Fernández R, Gabaldón T (2020) Gene gain and loss across the metazoan tree of life. Nature Ecology & Evolution 4:524

Fritzenwanker JH, Gerhart J, Freeman RM, Lowe CJ (2014) The Fox/Forkhead transcription factor family of the hemichordate Saccoglossus kowalevskii. EvoDevo 5:1

Gąsiorowski L, Chai C, Rozanski A, Purandare G, Ficze F, Mizi A, Wang B, Rink JC (2024) Regeneration in the absence of canonical neoblasts in an early branching flatworm. bioRxiv:2024.05.24.595708

Gąsiorowski L, Dittmann IL, Brand JN, Ruhwedel T, Möbius W, Egger B, Rink JC (2023a) Convergent evolution of the sensory pits in and within flatworms. BMC biology 21:1

Gąsiorowski L, Martín-Durán JM, Hejnol A (2023b) The evolution of Hox genes in Spiralia. In: Ferrier DEK (ed) Hox Modules in Evolution and Development. CRC Press, Boca Raton, pp. 177–194

Gregory TR, Hebert PDN, Kolasa J (2000) Evolutionary implications of the relationship between genome size and body size in flatworms and copepods. Heredity 84:201

Gross V, TreÑorn S, Reichelt J, Epple L, Lüter C, Mayer G (2019) Miniaturization of tardigrades (water bears): morphological and genomic perspectives. Arthropod structure & development 48:12

Hannenhalli S, Kaestner KH (2009) The evolution of Fox genes and their role in development and disease. Nature Reviews Genetics 10:233

Ivanković M, Brand JN, Pandolfini L, Brown T, Pippel M, Rozanski A, Schubert T, Grohme MA, Winkler S, Robledillo L, Zhang M, Codino A, Gustincich S, Vila-Farré M, Zhang S, Papantonis A, Marques A, Rink JC (2024) A comparative analysis of planarian genomes reveals regulatory conservation in the face of rapid structural divergence. Nature Communications 15:8215

Janssen R, Schomburg C, Prpic N-M, Budd GE (2022) A comprehensive study of arthropod and onychophoran Fox gene expression patterns. PLOS ONE 17:e0270790

Jennings J (1957) Studies on feeding, digestion, and food storage in free-living flatworms (Platyhelminthes: Turbellaria). The Biological Bulletin 112:63

Koinuma S, Umesono Y, Watanabe K, Agata K (2003) The expression of planarian brain factor homologs, DjFoxG and DjFoxD. Gene expression patterns 3:21

Kryuchkova-Mostacci N, Robinson-Rechavi M (2016) Tissue-Specificity of Gene Expression Diverges Slowly between Orthologs, and Rapidly between Paralogs. PLOS Computational Biology 12:e1005274

Kuehn E, Clausen DS, Null RW, Metzger BM, Willis AD, Özpolat BD (2022) Segment number threshold determines juvenile onset of germline cluster expansion in Platynereis dumerilii. Journal of Experimental Zoology Part B: Molecular and Developmental Evolution 338:225

Kumamoto T, Hanashima C (2017) Evolutionary conservation and conversion of Foxg1 function in brain development. Development, growth & differentiation 59:258

Langleib M, Calvelo J, Costábile A, Castillo E, Tort JF, Hoffmann FG, Protasio AV, Koziol U, Iriarte A (2024) Evolutionary analysis of genome-specific duplications in flatworm genomes. bioRxiv:2024.02.05.578899

Larroux C, Luke GN, Koopman P, Rokhsar DS, Shimeld SM, Degnan BM (2008) Genesis and Expansion of Metazoan Transcription Factor Gene Classes. Molecular Biology and Evolution 25:980

Larsson K, Jondelius U (2008) Phylogeny of Catenulida and support for Platyhelminthes. Organisms Diversity & Evolution 8:378

Laumer CE, Hejnol A, Giribet G (2015) Nuclear genomic signals of the ‘microturbellarian’ roots of platyhelminth evolutionary innovation. Elife 4

Lespinet O, Wolf YI, Koonin EV, Aravind L (2002) The role of lineage-specific gene family expansion in the evolution of eukaryotes. Genome research 12:1048

Lewin TD, Liao IJ-Y, Luo Y-J (2024) Conservation of animal genome structure is the exception not the rule. bioRxiv:2024.08.02.606322

Liao IJ-Y, Lu T-M, Chen M-E, Luo Y-J (2023) Spiralian genomics and the evolution of animal genome architecture. Briefings in Functional Genomics 22:498

Luo Y-J, Kanda M, Koyanagi R, Hisata K, Akiyama T, Sakamoto H, Sakamoto T, Satoh N (2018) Nemertean and phoronid genomes reveal lophotrochozoan evolution and the origin of bilaterian heads. Nature ecology & evolution 2:141

Malmstrøm M, Britz R, Matschiner M, Tørresen OK, Hadiaty RK, Yaakob N, Tan HH, Jakobsen KS, Salzburger W, Rüber L (2018) The most developmentally truncated fishes show extensive Hox gene loss and miniaturized genomes. Genome biology and evolution 10:1088

Mantica F, Iñiguez LP, Marquez Y, Permanyer J, Torres-Mendez A, Cruz J, Franch-Marro X, Tulenko F, Burguera D, Bertrand S, Doyle T, Nouzova M, Currie PD, Noriega FG, Escriva H, Arnone MI, Albertin CB, Wotton KR, Almudi I, Martin D, Irimia M (2024) Evolution of tissue-specific expression of ancestral genes across vertebrates and insects. Nature Ecology & Evolution 8:1140

Martín-Durán JM, Vellutini BC, Marlétaz F, Cetrangolo V, Cvetesic N, Thiel D, Henriet S, Grau-Bové X, Carrillo-Baltodano AM, Gu W, Kerbl A, Marquez Y, Bekkouche N, Chourrout D, Gómez-Skarmeta JL, Irimia M, Lenhard B, Worsaae K, Hejnol A (2021) Conservative route to genome compaction in a miniature annelid. Nature Ecology & Evolution 5:231

Mazet F, Yu J-K, Liberles DA, Holland LZ, Shimeld SM (2003) Phylogenetic relationships of the Fox (Forkhead) gene family in the Bilateria. Gene 316:79

Pascual-Carreras E, Herrera-Úbeda C, Rosselló M, Coronel-Córdoba P, Garcia-Fernàndez J, Saló E, Adell T (2021) Analysis of Fox genes in Schmidtea mediterranea reveals new families and a conserved role of Smed-foxO in controlling cell death. Scientific reports 11:2947

Plessy C, Mansfield MJ, Bliznina A, Masunaga A, West C, Tan Y, Liu AW, Grašič J, del Río Pisula MS, Sánchez-Serna G (2024) Extreme genome scrambling in marine planktonic Oikopleura dioica cryptic species. Genome Research 34:426

Price MN, Dehal PS, Arkin AP (2010) FastTree 2–approximately maximum-likelihood trees for large alignments. PloS one 5:e9490

Riddiford N, Olson PD (2011) Wnt gene loss in flatworms. Development genes and evolution 221:187

Rohde K (1991) The evolution of protonephridia of the Platyhelminthes. Hydrobiologia 227:315

Rohde K, Watson N (1993) Ultrastructure of the protonephridial system of regenerating Stenostomum sp.(Plathelminthes, Catenulida). Zoomorphology 113:61

Schindelin J, Arganda-Carreras I, Frise E, Kaynig V, Longair M, Pietzsch T, Preibisch S, Rueden C, Saalfeld S, Schmid B, Tinevez J-Y, White DJ, Hartenstein V, Eliceiri K, Tomancak P, Cardona A (2012) Fiji: an open-source platform for biological-image analysis. Nature Methods 9:676

Seudre O, Martín-Zamora FM, Rapisarda V, Luqman I, Carrillo-Baltodano AM, Martín-Durán JM (2022) The Fox gene repertoire in the annelid Owenia fusiformis reveals multiple expansions of the foxQ2 class in Spiralia. Genome Biology and Evolution 14:evac139

Shimeld SM, Degnan B, Luke GN (2010) Evolutionary genomics of the Fox genes: Origin of gene families and the ancestry of gene clusters. Genomics 95:256

Sievers F, Higgins DG (2014) Clustal Omega, Accurate Alignment of Very Large Numbers of Sequences. In: Russell DJ (ed) Multiple Sequence Alignment Methods. Humana Press, Totowa, NJ, pp. 105–116

Slyusarev GS, Starunov VV, Bondarenko AS, Zorina NA, Bondarenko NI (2020) Extreme Genome and Nervous System Streamlining in the Invertebrate Parasite <em>Intoshia variabili</em>. Current Biology 30:1292

Stentiford GD, Dunn AM (2014) Microsporidia in aquatic invertebrates. Microsporidia: Pathogens of opportunity:579

Stubbs JL, Oishi I, Izpisua Belmonte JC, Kintner C (2008) The forkhead protein Foxj1 specifies node-like cilia in Xenopus and zebrafish embryos. Nature genetics 40:1454

Susumu O (1970) Evolution by Gene Duplication. Allen and Unwin, London

Toresson H, Martinez-Barbera JP, Bardsley A, Caubit X, Krauss S (1998) Conservation of BF-1 expression in amphioxus and zebrafish suggests evolutionary ancestry of anterior cell types that contribute to the vertebrate telencephalon. Development genes and evolution 208:431

Tyler S (1984) Turbellarian Platyhelminths. In: Bereiter-Hahn J, Matoltsy AG, Richards KS (eds) Biology of the Integument: Invertebrates. Springer Berlin Heidelberg, Berlin, Heidelberg, pp. 112–131

Vij S, Rink JC, Ho HK, Babu D, Eitel M, Narasimhan V, Tiku V, Westbrook J, Schierwater B, Roy S (2012) Evolutionarily Ancient Association of the FoxJ1 Transcription Factor with the Motile Ciliogenic Program. PLOS Genetics 8:e1003019

Vu HT-K, Mansour S, Kücken M, Blasse C, Basquin C, Azimzadeh J, Myers EW, Brusch L, Rink JC (2019) Dynamic polarization of the multiciliated planarian epidermis between body plan landmarks. Developmental cell 51:526

Vu HT-K, Rink JC, McKinney SA, McClain M, Lakshmanaperumal N, Alexander R, Sánchez Alvarado A (2015) Stem cells and fluid flow drive cyst formation in an invertebrate excretory organ. Elife 4:e07405

Wiberg RAW, Brand JN, Viktorin G, Mitchell JO, Beisel C, Schärer L (2023) Genome assemblies of the simultaneously hermaphroditic flatworms Macrostomum cliftonense and Macrostomum hystrix. G3 Genes|Genomes|Genetics 13

Wolf FA, Angerer P, Theis FJ (2018) SCANPY: large-scale single-cell gene expression data analysis. Genome Biology 19:15

Yu X, Ng CP, Habacher H, Roy S (2008) Foxj1 transcription factors are master regulators of the motile ciliogenic program. Nature genetics 40:1445

Zadesenets KS, Ershov NI, Berezikov E, Rubtsov NB (2017a) Chromosome Evolution in the Free-Living Flatworms: First Evidence of Intrachromosomal Rearrangements in Karyotype Evolution of Macrostomum lignano (Platyhelminthes, Macrostomida). Genes 8:298

Zadesenets KS, Schärer L, Rubtsov NB (2017b) New insights into the karyotype evolution of the free-living flatworm Macrostomum lignano (Platyhelminthes, Turbellaria). Scientific Reports 7:6066

