## Supplementary Figures S1-S15 for "Evidence for multiple independent expansions of Fox gene families within flatworms"

*Journal of Molecular Evolution*

Ludwik Gąsiorowski [1,*]

[1] Institute of Evolutionary Biology, Faculty of Biology University of Warsaw, ul. Żwirki i Wigury 101, 02-089 Warsaw, Poland; ORCID 0000-0003-2238-7587

**Online Resource 3:**

Supplementary Figures S1-S15

**Fig. S1.** Phylogeny of metazoan Fox genes, approximately-maximum-likelihood tree computed with FastTree (v 2.1.11) under LG amino acid substitution model. Flatworm sequences are in red. Reference sequences come from Fritzenwanker et al. 2014 and NCBI database (listed in Online Resource: Table S2

**Fig. S1.** Phylogeny of metazoan Fox genes, maximum-likelihood tree computed with raxmlGUI (v 2.0) under LG amino acid substitution model. Flatworm sequences are in red. Reference sequences same as in Fig. S1.

**Figs. S2-S15.** Phylogenies of particular Fox families in flatworms. Sequences of *Stenostomum leucops*, *Stenostomum brevipharyngium*, *Macrostomum cliftonense* and *Macrostomum hystrix* come from this study, the other sequences come from Pascual-Carreras et al. 2021 and are also listed as Online Resource 2. Reconstructed duplications of Fox genes are indicated with red arrows.
